# Multi-omic characterization of axolotl perilymph-cerebrospinal fluid reveals shifts in composition during limb regeneration

**DOI:** 10.64898/2026.08.27.747356

**Authors:** Noah Lopez, Bingsen Zhang, Steven R. Shuken, Yudong Zhou, Duygu Payzin-Dogru, Julia C. Paoli, Anthony E. Striker, S. Y. Celeste Wu, Tia Patel, Karly Chan, Sebastian Böhm, Hani D. Singer, Ashley R. Juarez, Ryan T. Kim, Laurel Shugart, Edward T. Chouchani, Jessica L. Whited

## Abstract

The axolotl salamander can fully regenerate amputated limbs, yet the systemic consequences underlying this process remain largely understudied. Cerebrospinal fluid is an emerging signaling medium capable of communicating with both the central and peripheral nervous systems, but its composition and potential role in salamander limb regeneration have not yet been examined using modern multi-omics techniques. Here, we developed a protocol for extracting mixed perilymph-cerebrospinal fluid (P-CSF) from axolotl and provided the first proteomic and metabolomic characterization of this biological fluid. We identified 2,626 unique proteins and 173 high-confidence metabolites and quantified them across four time points of early limb regeneration. We demonstrated that limb amputation drives progressive shifts in P-CSF proteins, including an elevation of sarcomeric muscle proteins, regeneration-associated factors, and protease/extracellular matrix proteins. We observed shifts in metabolites involved in oxidative stress, polyunsaturated fatty acid oxidation, and histamine metabolism. Injury-comparison experiments revealed that the observed proteomic changes as a result of limb amputation are different than crush injury, denervation, or tail amputation. This study proposes axolotl P-CSF as a reservoir for limb amputation-associated systemic signaling and as a potential conduit of signals involved in limb regeneration.

## Introduction

Accomplishing the regeneration of complex tissues has long been a goal of medical science. Humans are incapable of regenerating lost appendages, such as full limbs^1^. Furthermore, many of those who suffer from limb loss exhibit enhanced hypersensitization or suffer from phantom-limb pain (PLP)^2,3^. Elucidating how traumatic injury changes systemic physiology, including central nervous system (CNS) activity, could accelerate the development of treatments for many of the adverse health outcomes associated with limb amputation^4^. Understanding how these features change when a limb is successfully regenerated, for example in a species that regenerates full limbs, may provide critical comparative insights into such mechanisms.

The axolotl salamander is a vertebrate capable of regenerating full limbs^5^, and for which many modern experimental and genomic technologies have now been developed^6^. The formation of a blastema is required for axolotl limb regeneration. At 3 days post amputation (dpa), rapid reepithelialization has occurred and the wound epidermis has formed. By 7dpa, the apical epithelial cap has formed and begun recruiting blastema cells. At 14dpa, the blastema is fully established, and blastema markers such as kazal-type serine peptidase inhibitor domain 2 (KAZALD2) are robustly expressed^7^. Evaluating systems-level responses to limb amputation during these critical stages can further our understanding of how regeneration-competent organisms respond to major trauma and propagate pro-regenerative responses.

In recent years, there has been an expansion in the known functional roles of cerebrospinal fluid (CSF)^8,9^. CSF is a clear, colorless fluid that occupies the ventricles and subarachnoid space of the CNS. In mammals, CSF is mainly produced by the choroid plexus and is largely composed of water, ions, lipids, proteins, and contains small numbers of immune cells^10^. The blood-CSF barrier (BCSFB), primarily comprised of choroid plexus epithelial cells and the tight junctions between them, typically regulates the selective passage of water, electrolytes, and certain circulating plasma proteins into the CSF^11^. In models of oxidative stress^12^, aging^13^, or neuroinflammation^14^, this barrier can become compromised and can have profound effects on brain homeostasis. A potential mediator of this process is the activity of serine proteases and similar proteases^15^, whose function have also been associated with modulating growth factors and extracellular matrix (ECM) during salamander limb regeneration^16,17^.

While the historical understanding of CSF function centered on mechanical protection and waste removal from the CNS, a renewed interest in the role of CSF factors as neuromodulators and as a source of disease biomarkers has recently emerged^18–20^. We hypothesized that components of the CSF change during axolotl limb amputation and may regulate or coincide with aspects of the regenerative response. As a first step towards testing this hypothesis, we sought to create a foundational resource by providing the first proteomic and metabolomic characterization of axolotl CSF before and after limb amputation.

## Results

### Mixed perilymph-cerebrospinal fluid (P-CSF) is a distinct biofluid from blood

Our first goal was to develop a surgical procedure for extracting axolotl CSF. The presence of blood is a common complication of CSF collection and has significant consequences on downstream analyses^21^. The axolotl cranial cavity contains a complex network of brain vasculature^22^, making the extraction of clean CSF without the disruption of these fragile structures technically challenging. To avoid blood contamination, we used the otic capsule as a collection route (Figure 1A). The otic capsule is relatively spacious and has fewer blood vessels compared to the cranial cavity, making it a promising candidate for blood-free collection. We hypothesized that harvesting mixed perilymph-CSF (P-CSF) through this route would provide a clean biofluid for study while simultaneously serving as a broad assessment of the neural environment, similar to what has been proposed in humans^23^.

**Figure 1:**
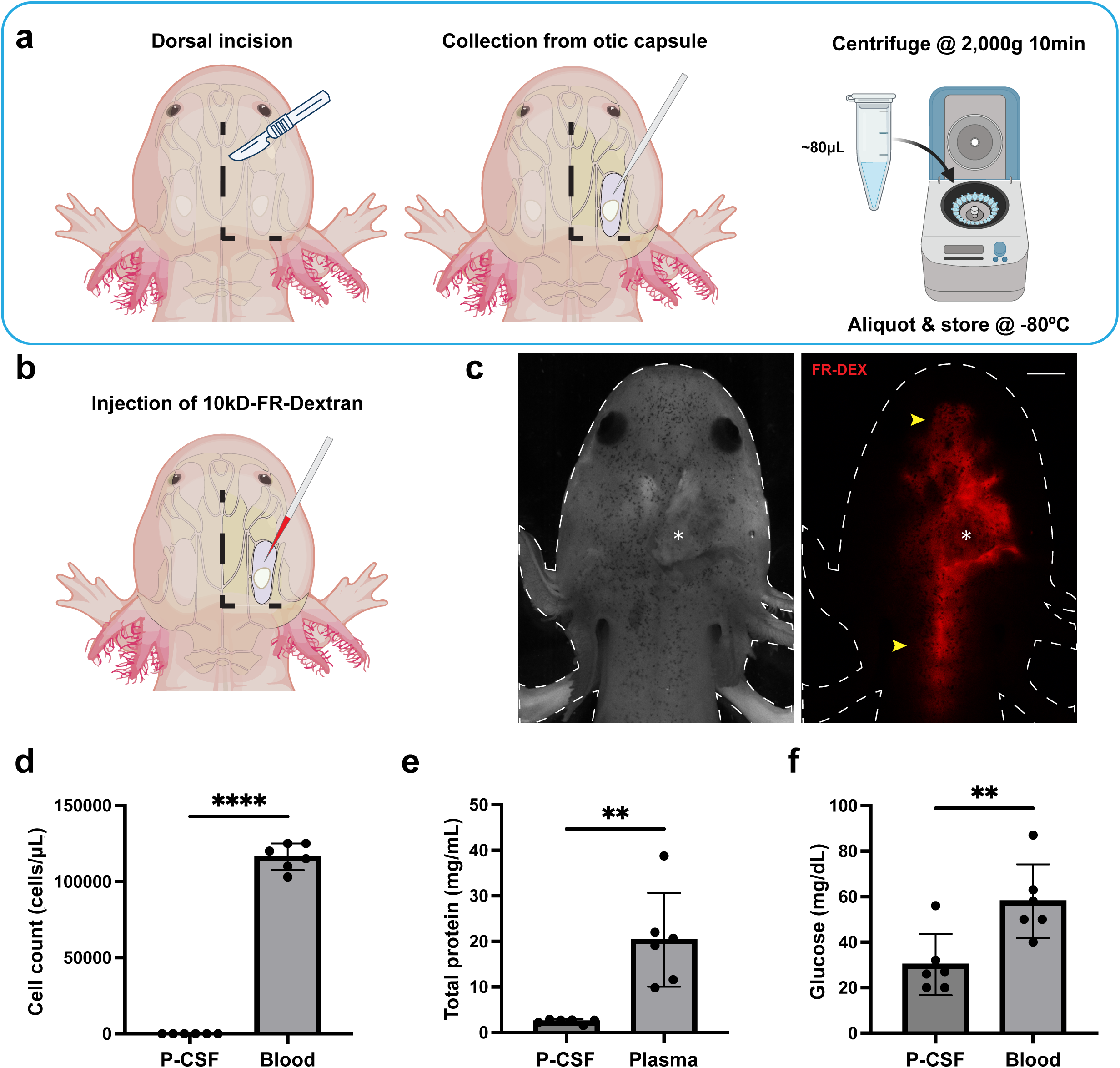
Establishing P-CSF as a distinct biological fluid. **a** Schematic of P-CSF extraction procedure. **b** Schematic of injection of 10 µl of 10-kDa Fluoro Ruby-conjugated dextran into the otic capsule **c** Images of an axolotl injected with 10 µl of 10-kDa Fluoro Ruby-conjugated dextran into the otic capsule. The left image was taken in brightfield and was converted to grayscale. The right image was taken in the red channel. White asterisk (*) indicates the injection site and typical P-CSF collection site. Yellow arrows (→) represent diffusion of the dextran into the cranial cavity and spinal cord. Scale bar = 2 mm. **d** Mean cell counts in intact P-CSF and whole blood. Two-sided unpaired t-test with Welch’s correction (p-value = < 0.0001). Data are presented as mean ± s.d. ****: p < 0.0001. **e** Mean total protein concentration of intact P-CSF vs EDTA plasma using BCA. All samples ran in triplicate and averages represented. Two-sided unpaired t-test with Welch’s correction (p-value = 0.0081). Data are presented as mean ± s.d. **: p < 0.01. **f** Mean glucose concentration of intact P-CSF and whole blood using a glucose meter. Two-sided unpaired t-test with Welch’s correction (p-value = 0.0094). Data are presented as mean ± s.d. **: p < 0.01.

Using this approach, we successfully extracted an average of ∼80µL clear P-CSF per animal. To confirm the continuity between the otic capsule and the cranial cavity, we injected fluorescently labeled 10-kDa dextran into the collection site (Figure 1B). The injected dye rapidly diffused into the cranial cavity and spinal cord, demonstrating direct continuity between the extracted compartment and the CSF-containing cranial cavity (Figure 1C). We then assessed sample quality using multiple complementary metrics to demonstrate the absence of blood. Any samples containing visible erythrocytes before or after centrifugation were excluded. The mean total cell count of P-CSF was 10 ± 5 cells/µL, compared to 117,500 ± 8,756 cells/µL in whole blood (Figure 1D). Mean total protein concentration of P-CSF was 2.5 ± 0.5 mg/mL, compared to 20.3 ± 10.3 mg/mL in EDTA blood plasma (Figure 1E). Mean P-CSF glucose was 30.2 ± 13.5 mg/dL compared to 58.0 ± 16.2 mg/dL in whole-blood (Figure 1F). Together, these results confirm the technical viability of the otic capsule extraction procedure and establish the biological distinction between P-CSF and peripheral blood.

### High-resolution quantitative mass spectrometry with tandem mass tags reveals a diverse protein repertoire in axolotl P-CSF

To assess whether limb amputation alters the repertoire of proteins in axolotl P-CSF, we performed TMT labeling followed by HPLC-based fractionation to quantify proteins at maximal depth with high quantitative accuracy and precision^24^. We profiled P-CSF in homeostatic intact animals and at three time points in early limb regeneration (Figure 2A). No significant differences in total protein concentration or hemoglobin were observed between groups (Supplementary Fig. 1A/B). Combining all time points, we identified 2,626 unique proteins in axolotl P-CSF (Supplementary Fig. 1C) (Supplementary Table 1). 87.5% of identified proteins had two or more unique peptides (Supplementary Fig. 1D). Peptides were matched to the axolotl reference proteome (NCBI RefSeq Assembly). The axolotl genome annotation assigns placeholder identifiers (e.g., LOCXXXXXXXXX) to uncharacterized or computationally predicted loci^25^. To make these genes compatible with downstream analyses, LOC-annotated entries were assigned the closest matching human ortholog gene symbol using the NCBI gene description where applicable. Proteins sharing the same gene symbol were distinguished by sequentially appending a numeric suffix (e.g., GENE_1, GENE_2). It should be noted that while these specific genes may resemble human orthologs, their particular function or expression profiles may not be fully equivalent in axolotl.

**Figure 2:**
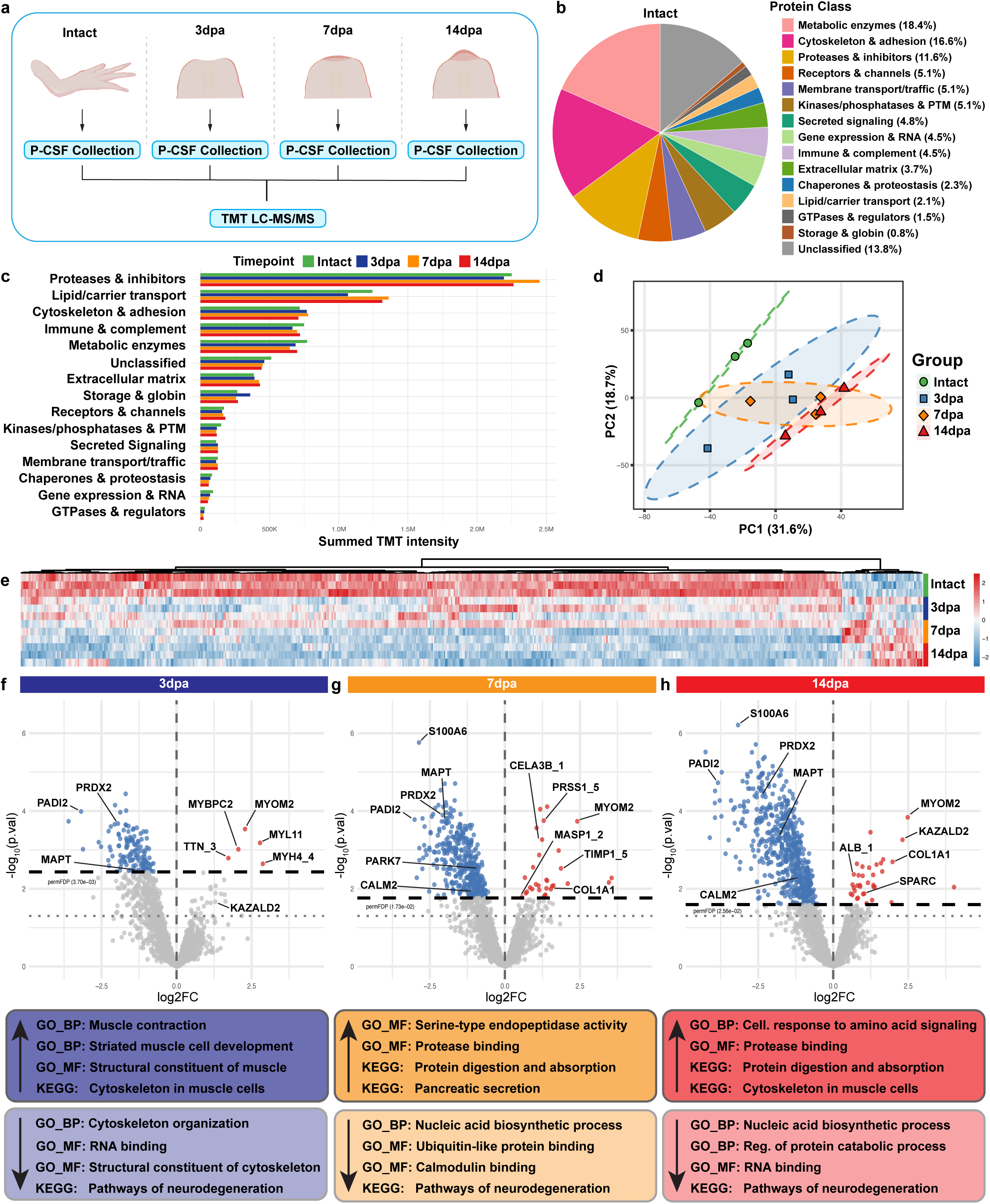
High resolution TMT-labeled quantitative mass spectrometry proteomics of P-CSF from different timepoints of limb amputation. **a** Schematic of P-CSF sample groups. Animals were either left intact or amputated on all four limbs. Each sample represents individual animals. **b** Protein class pie chart of proteins found in P-CSF. Each percentage indicated the proportion of all identified proteins by name. **c** Protein class representation by relative abundance. Relative protein abundance is represented on the x-axis. Protein class is represented on the y-axis. **d** PCA plot of P-CSF limb amputation time points. **e** Heatmap of differentially abundant proteins in P-CSF after limb amputation. **f-h** Volcano plots for Intact vs 3dpa, Intact vs 7dpa, or Intact vs 14dpa P-CSF proteins. Red dots indicate significantly elevated. Blue dots indicate significantly decreased. Grey dotted line represents nominal p-value significance threshold (p = 0.05). Black dotted line permFDP adjusted threshold for significance. permFDP significance thresholds for Intact vs 3dpa (permFDP < 3.70e-03), Intact vs 7dpa (permFDP < 1.73e-02), and Intact vs 14dpa (permFDP < 2.56e-02). Colored boxes contain gene ontology (GO) and KEGG representative terms for each comparison.

The most consistently detected proteins across intact P-CSF samples were serum albumin-like (LOC138515996; ALB_2), complement C3 (C3), serotransferrin-A-like (LOC138491525; TF_2), apolipoprotein B-100 (APOB), and alpha-2-macroglobulin-like protein 1 (LOC138580212; A2ML1_2). Each protein was assigned a protein class using the PANTHER classification system^26^ and then manually consolidated into generalized categories (Supplementary Table 2). By number of identified proteins belonging to each class, the most represented classes were metabolic enzymes (18.4%), cytoskeletal and adhesion proteins (16.6%), unclassified proteins (13.8%), and proteases & inhibitors (11.6%) (Figure 2B). When ranked by summed TMT intensity, proteases & inhibitors and lipid/carrier transport proteins emerged as the dominant classes (Figure 2C). The observed disparity between represented protein class members and summed TMT intensities suggests that while metabolic enzymes may contribute to a broad functional repertoire in axolotl P-CSF, proteases/inhibitors and lipid transport proteins may represent a readily detectable and substantial portion of P-CSF proteins. This is consistent with the known biochemical characteristics of perilymph and CSF as a protease-regulated, lipid-rich signaling fluid^27,28^.

### Limb amputation results in elevated muscle components and regeneration-associated proteins in P-CSF

We next characterized how the protein repertoire of P-CSF changes in response to limb amputation and during the initiation of regeneration. Principal component analysis (PCA) of samples showed visual clustering between time points, with intact and 14dpa groups being the most spatially distinct from one another (Figure 2D). PC1 explained 31.6% and PC2 explained 18.7% of total variance. Individual PCA loadings can be referred to in (Supplementary Table 3). Differential relative protein abundance testing identified 5 upregulated and 134 downregulated proteins at 3dpa, 30 upregulated and 376 downregulated at 7dpa, and 35 upregulated and 545 downregulated at 14dpa (Figures 2E-H). GO and KEGG pathway enrichment was performed on each direction and time point (Supplementary Tables 4–5).

At 3dpa, upregulated proteins were enriched for terms such as muscle contraction and striated muscle tissue development. This was driven by sarcomeric proteins including myosin-4-like (LOC138582820; MYH4) (log2FC = 2.87), myosin regulatory light chain 11 (MYL11) (log2FC = 2.78), myomesin-2 (MYOM2) (log2FC = 2.28), and titin (TTN) (log2FC = 1.73). These proteins and related orthologs are broadly conserved as muscle cell components^29^ and have been detected in human CSF in healthy individuals^30^. This pattern persisted at 7dpa, with continued enrichment for structural constituents of muscle and muscle tissue morphogenesis, represented by MYOM2 (log2FC = 2.42), telethonin-like (LOC138521710; TCAP) (log2FC = 2.32), and TTN (log2FC = 1.41).

At 7dpa, we also observed an increase in serine proteases and ECM-related proteins. GO terms emerged including serine-type endopeptidase activity, protease binding, and KEGG terms for protein digestion and absorption. These corresponded to proteins including collagen alpha-1 chain (COL1A1) (log2FC = 1.57), chymotrypsin-like protease CTRL-1 (LOC138447663; CTRL_1) (log2FC = 1.41), trypsin-like (LOC138580108; PRSS1_5) (log2FC = 1.29), and proproteinase E-like (LOC138472660; CELA3B_1) (log2FC = 1.24). A slight increase in mannan-binding lectin serine protease 1 (MASP1_2) (log2FC = 0.56) was also observed. Interestingly, we observed an elevation of VWDE (log2FC = 1.28), a known feature of axolotl limb regeneration^31^. Notable increases in inner ear-associated proteins including otogelin (OTOG) (log2FC = 3.56) and otoconin-90 (OC90) (log2FC = 2.09) were also observed. The observed changes in relative protein abundance of ear-associated proteins may hint at either a local otic response or varying ratios of perilymph to CSF as a consequence of the extraction procedure. However, it should be noted that OTOG has also been shown to be expressed in the axolotl blastema by small secretory cells of the epithelium during limb regeneration^32^.

At 14dpa, enriched terms included extracellular matrix structural constituent conferring tensile strength and protease binding. These were driven by multiple collagens: COL1A1 (log2FC = 1.98), COL1A2 (log2FC = 1.63), COL6A1 (log2FC = 0.89), and COL11A1 (log2FC = 1.11). MYOM2 (log2FC = 2.48) and TCAP (log2FC = 1.66) remained elevated. Serum albumin-like (LOC138515210; ALB_1) (log2FC = 0.97) was also increased, which may be relevant given that an elevated CSF/blood albumin ratio is a recognized clinical indicator of blood-brain barrier (BBB) perturbation^33^. Strikingly, two established blastema-associated proteins, kazal-type serine protease inhibitor domain-containing protein 2-like (LOC138516914; KAZALD2) (log2FC = 2.31) and secreted protein acidic and cysteine rich (SPARC) (log2FC = 1.03), were upregulated at 14dpa. Both are known to be upregulated during axolotl limb regeneration and are actively secreted by the blastema^32,34^. These results raised the question of whether elevated P-CSF proteins after amputation reflect a systemic injury response signature, local blastema-derived secretion reaching the CSF compartment, or a combination of the two.

### Trajectory analysis reveals distinct temporal trends in P-CSF after limb amputation

The temporal structure of our proteomics dataset enabled trajectory-based clustering analysis, grouping proteins by shared temporal patterns in normalized intensity across time rather than by significance at individual contrasts. This approach complements contrast-level differential analysis by capturing proteins whose dynamics emerge as a coherent pattern across the regenerative time course, even those whose comparisons at any single time point may fall below conventional significance thresholds. Using this, we identified six distinct temporal trajectories in P-CSF after limb amputation (Figure 3A, 3B).

**Figure 3:**
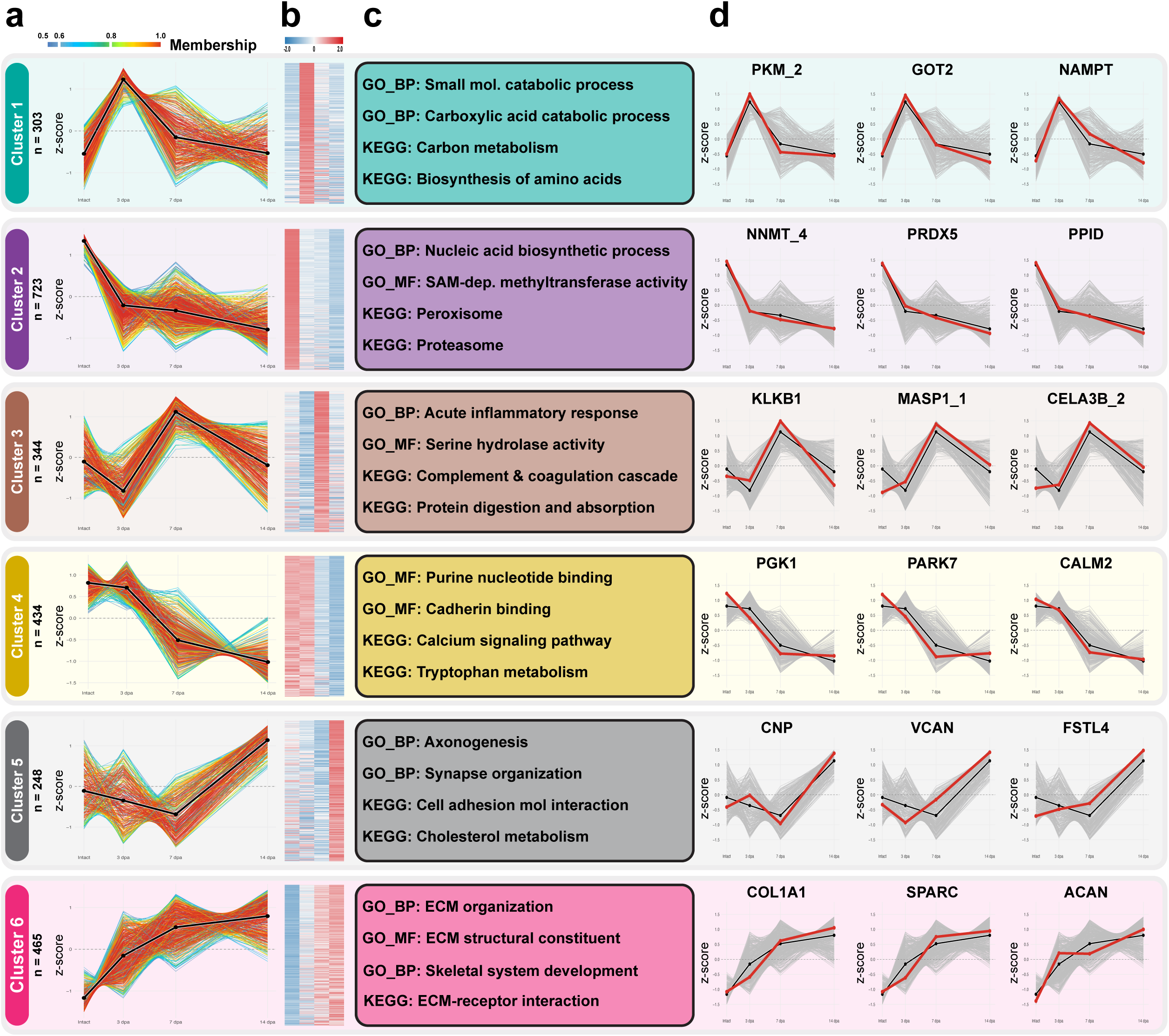
Mfuzz trajectory clustering of P-CSF limb amputation timeseries proteomics. **a** Six clusters with distinct temporal trajectories. Protein membership counts are indicated for each cluster. Trajectory plots are depicted for each cluster. Black line indicates the cluster centroid. Colored lines indicate fuzzy membership score (0-1 scale; warm colors indicate high membership). **b** Heatmap of z-scored protein abundance for all clustered proteins, separated by individual cluster assignments. **c** Representative GO and KEGG pathway enrichments for each cluster. **d** Individual trajectory plots for three representative proteins per cluster (black line = cluster mean, red line = individual protein trajectory, gray lines = all other protein trajectories in cluster).

Cluster 1 (n = 303 proteins), representing a transient “immediate injury response”, is characterized by a rapid increase in relative protein abundance at 3dpa followed by a gradual return toward baseline. Relevant GO/KEGG terms for this cluster included small molecule catabolic processes, carbon metabolism, and biosynthesis of amino acids (Figure 3C). Example cluster members include pyruvate kinase (PKM), aspartate aminotransferase, mitochondrial (GOT2), and nicotinamide phosphoribosyltransferase (NAMPT) (Figure 3D). Cluster 2 (n = 723 proteins) represents the largest cluster and describes a persistent “immediate-suppressed” trajectory. This cluster is characterized by a sustained decrease beginning at 3dpa that continues through 14dpa. Relevant GO/KEGG terms were nucleic acid biosynthetic process, SAM-dependent methyltransferase activity, peroxisome, and proteasome. Example cluster members include nicotinamide N-methyltransferase-like (LOC138487683; NNMT_2), peroxiredoxin-5 (PRDX5), and peptidyl-prolyl cis-trans isomerase D (PPID). Cluster 3 (n = 344 proteins) defines a “mid-stage protease-associated” trajectory, with relative protein abundance peaking at 7dpa before returning toward baseline. Enriched terms included acute inflammatory response, serine protease activity, and complement and coagulation cascade. Representative members include plasma kallikrein-like (LOC138516427; KLKB1), mannan-binding lectin serine protease 1 (MASP1_1), and proproteinase E-like (LOC138472662; CELA3B_2), suggesting a window of active protease activity. Cluster 4 (n = 434 proteins), a “mid-stage suppression” trajectory, is characterized by a sustained decrease at 7dpa and 14dpa. In contrast to Cluster 2, which declines immediately at 3dpa, this cluster’s later onset suggests a secondary suppressive wave that coincides with active blastema formation. Enriched terms included purine nucleotide binding, cadherin binding, and calcium signaling. Representative members include Parkinson disease protein 7 (PARK7), phosphoglycerate kinase 1 (PGK1), and calmodulin-2 (CALM2). Cluster 5 (n = 248 proteins) defines a “late-peak” trajectory, with a distinct increase in relative abundance at 14dpa. Enriched terms included axonogenesis, synapse organization, and cell adhesion molecule interaction. Representative members include 2’,3’-cyclic-nucleotide 3’-phosphodiesterase (CNP), versican core protein-like (VCAN), and follistatin-related protein 4 (FSTL4) that may be associated with nerve-associated processes or neural turnover in the P-CSF compartment or altered ECM reorganization^35–37^. Finally, Cluster 6 (n = 465 proteins) describes a “gradual increase” trajectory, with relative protein rising steadily across all post-amputation time points. Enriched terms included ECM organization, ECM structural constituent, and skeletal system development. These terms may be associated with progressive connective tissue remodeling and skeletal patterning occurring during limb regeneration. Representative members include COL1A1, SPARC, and aggrecan core protein (ACAN). Individual cluster membership and associated GO/KEGG terms for all clusters can be found in Supplementary Tables 6-8.

### Limb amputation elicits a distinct P-CSF proteomics response

To test whether the observed changes in relative P-CSF protein abundance were unique to limb amputation and blastema formation, we compared three alternative injury models: limb denervation, limb crush, and distal tail amputation (Figure 4A). This new cohort was collected at 14 days post injury to mirror the time point in which limb regeneration-associated proteins were most prominent. No change in total P-CSF protein concentration was observed in the injury groups relative to intact (Supplementary Fig. 2A). While a statistically significant difference in hemoglobin concentration was observed between the denervation and tail amputation group compared to the intact condition (Supplementary Fig. 2B), the absolute difference was small, and all values remained within an acceptable range for downstream proteomic analysis. Dynamic range of relative protein abundance showed comparable protein distributions across groups, though only 2,209 proteins were recovered in this cohort (Supplementary Fig. 2C) (Supplementary Table 9). PCA showed spatial clustering between injury models. 28.5% of the variability can be explained by PC1 and 16.6% of the variability can be explained by PC2 (Figure 4B). Individual PCA loadings can be referred to in (Supplementary Table 10).

**Figure 4:**
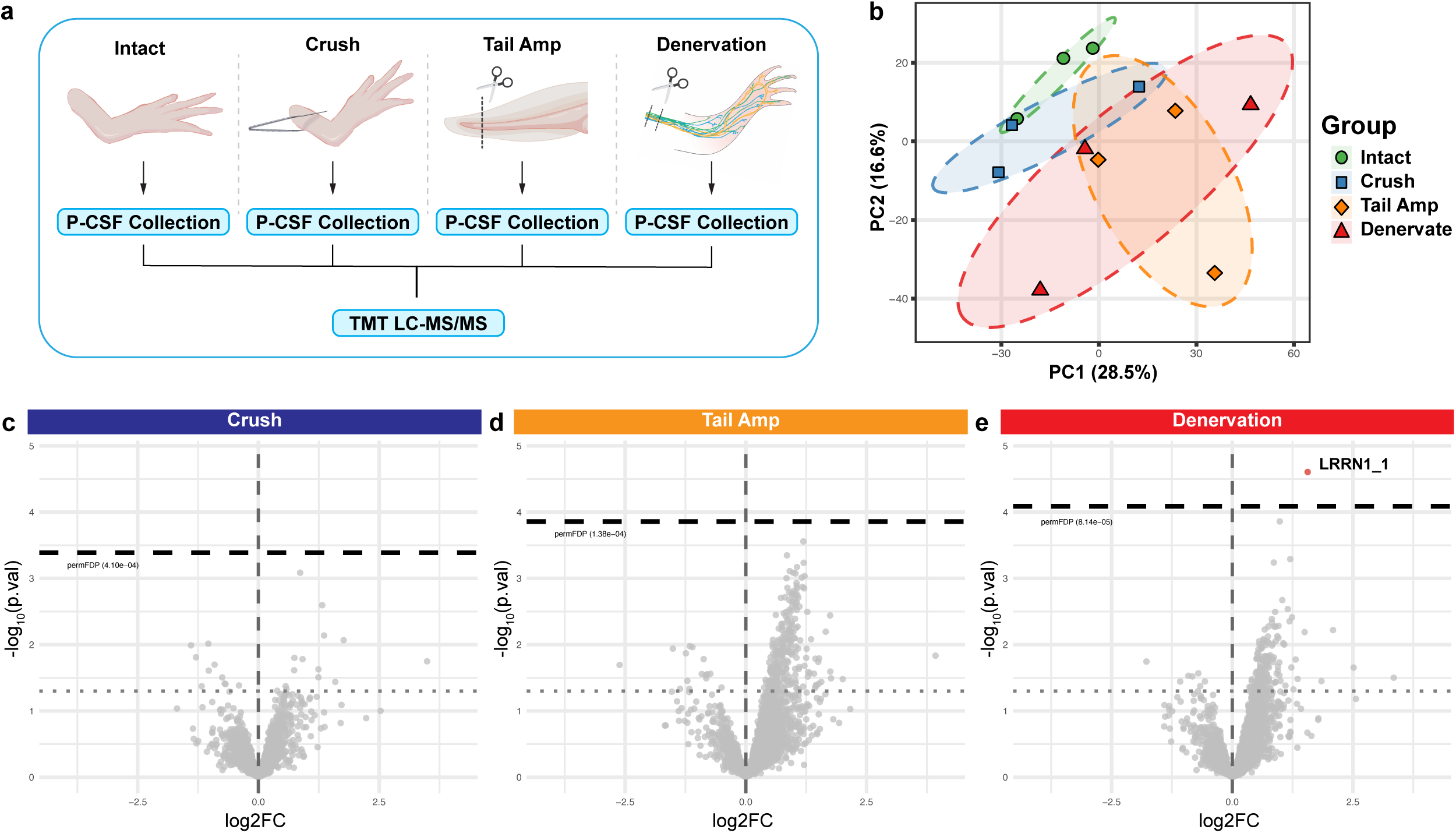
High resolution TMT-labeled quantitative mass spectrometry proteomics of P-CSF from different injury models. **a** Schematic of P-CSF injury model groups. **b** PCA plot of P-CSF injury groups. **c-e** Volcano plots for Intact vs Crush, Intact vs Tail Amputation, or Intact vs Denervation P-CSF proteins. Red dots indicate significantly elevated. Blue dots indicate significantly decreased. Grey dotted line represents nominal p-value significance threshold (p = 0.05). Black dotted line permFDP adjusted threshold for significance. permFDP significance thresholds for Intact vs Crush (permFDP < 4.10e-04), Intact vs Tail Amputation (permFDP < 1.38e-04), and Intact vs Denervation (permFDP < 8.14e-05).

Differential relative protein abundance testing revealed that only one protein in the denervation group, leucine-rich repeat neuronal protein 1-like (LOC138569445; LRRN1_2) (log2FC = 1.03), was significantly differentially abundant relative to intact P-CSF. LRRN1 was also significantly elevated at 14dpa (log2FC = 0.80) and is a transmembrane protein with mammalian expression enriched in the CNS and, notably in cochlear sensory epithelium^38,39^. LRRN1 elevation in the denervation group may suggest a systemic response to limb nerve transection or potentially reflect a local inner ear response. No significant differentially abundant proteins were identified in the crush and tail amputation groups (Figure 4C-E). Under these conditions, the proteomic changes observed following limb amputation were not broadly recapitulated by the alternate injury models at the examined timepoints and may require further testing of injury models or timepoints.

### Untargeted metabolomics reveals a biochemically diverse small-molecule repertoire in axolotl P-CSF

To complement the proteomics and provide a characterization of small-molecule composition, we profiled the P-CSF metabolome across the same four limb amputation time points (Figure 5A). Untargeted metabolomics was performed using hydrophilic interaction liquid chromatography coupled to mass spectrometry (HILIC LC-MS), a platform well-suited for the detection of polar metabolites and polyamine derivatives in biological samples^40,41^. After filtering for annotation quality, inspecting elution profiles, and considering compound origin, we detected and quantified 173 level-2 endogenous metabolites (Supplementary Table 11). The most consistently detected metabolites across intact P-CSF samples were betaine, creatine, lactic acid, hypoxanthine, and pyroglutamic acid. Together, these metabolites reflect one-carbon cycling^42^, energy metabolism^43^, and purine turnover^44^ as prominent features of the homeostatic P-CSF chemical environment.

**Figure 5:**
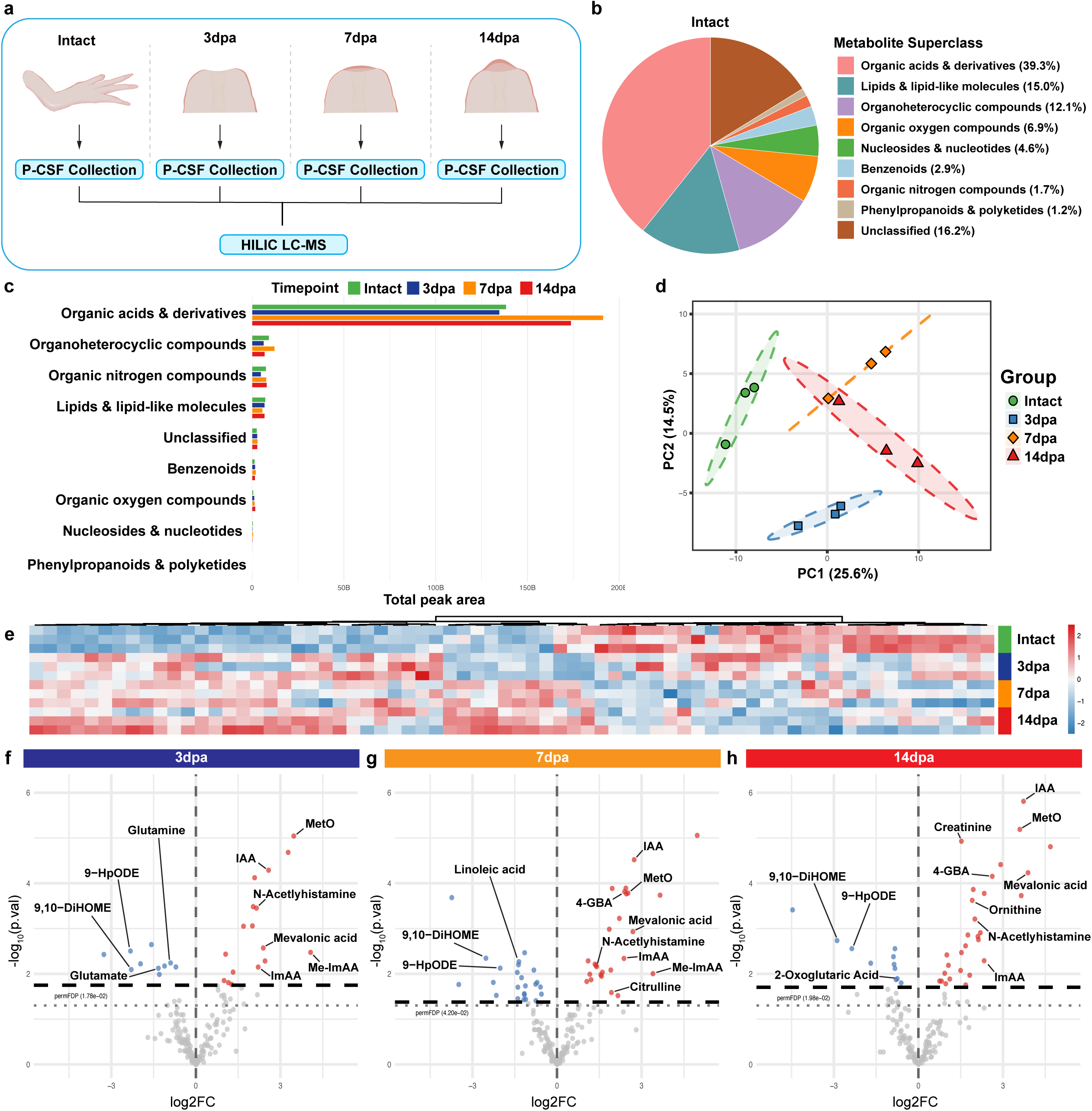
Untargeted metabolomics of P-CSF from different timepoints of limb amputation. **a** Schematic of P-CSF sample groups. Animals were either left intact or amputated on all four limbs. Each sample represents individual animals. **b** Metabolite superclass pie chart of metabolites found in P-CSF. Each percentage indicates the proportion of all identified metabolites by name. **c** Metabolite superclass representation by summed peak area. Total peak area represented on the x-axis. Metabolite superclass represented on the y-axis. **d** PCA plot of P-CSF metabolite timepoints. **e** Heatmap of differentially abundant metabolites in P-CSF after limb amputation. **f-h** Volcano plots for Intact vs 3dpa, Intact vs 7dpa, or Intact vs 14dpa P-CSF metabolites. Red dots indicate significantly elevated. Blue dots indicate significantly decreased. Grey dotted line represents nominal p-value significance threshold (p = 0.05). Black dotted line permFDP adjusted threshold for significance. permFDP significance thresholds for Intact vs 3dpa (permFDP < 1.78e-02), Intact vs 7dpa (permFDP < 4.20e-02), and Intact vs 14dpa (permFDP < 1.98e-02). Colored boxes contain gene ontology (GO) and KEGG representative terms for each comparison.

Using the HMDB metabolite classification system^45^, the detected P-CSF metabolites spanned 8 superclasses, with organic acids & derivatives being the largest represented fraction (39.3%), lipids & lipid-like molecules (15.0%), organoheterocyclic compounds (12.1%), organic oxygen compounds (6.9%%), nucleosides & nucleotides (4.6%), benzenoids (2.9%), organic nitrogen compounds (1.7%), and phenylpropanoids & polyketides (1.2%). All metabolites without clear assignment were grouped into unclassified (16.2%) (Figure 5B). When superclasses were ranked by total summed detection intensity, organic acids & derivatives were the most consistently identified metabolites in our dataset (Figure 5C). This superclass can be further broken down into subclasses and primarily consisted of amino acids & derivatives, carboxylic acid derivatives, dicarboxylic acid derivatives, and tricarboxylic acid derivatives.

### Limb amputation results in changes to P-CSF metabolites

Consistent with the proteomics, PCA of P-CSF metabolite profiles showed that intact samples clustered distinctly from the amputated conditions. PC1 and PC2 explained 25.6% and 14.5% of total variance, respectively (Figure 5D). Individual PCA loadings can be referred to in (Supplementary Table 12). Differential relative abundance testing identified 17 increased and 10 decreased metabolites at 3dpa, 26 increased and 24 decreased at 7dpa, and 29 increased and 11 decreased at 14dpa (Figures 5E-H). A transient decrease in glutamate and glutamine was observed at 3dpa, suggesting perturbation of amino acid homeostasis at the earliest post-amputation time point.

Several metabolites showed consistent directional changes across all amputation time points. Among persistently increased metabolites, methionine sulfoxide (MetO), a stable oxidation product of methionine and a sensitive marker of ROS activity^46^, was elevated at all three time points. Indole-3-acetic acid (IAA), a tryptophan-derived metabolite associated with gut microbiome activity^47^, was also consistently elevated. This raises the possibility that amputation-associated physiological changes may influence microbiome-derived signaling in the P-CSF compartment or that disruptions in BCSFB permeability leads to the altered presence of these metabolites. However, the production of endogenous host-derived IAA cannot be explicitly ruled out^48^.

Mevalonic acid, a key intermediate in the mevalonate pathway and a precursor to cholesterol and isoprenoid biosynthesis^49^, was increased at all time points. Increases in histamine-related metabolites including N-acetylhistamine^50^, imidazoleacetic acid (ImAA)^51^, and methylimidazoleacetic acid (Me-ImAA)^52^ were observed across multiple time points, suggesting altered histaminergic metabolism as a consistent feature of the post-amputation P-CSF environment.

We observed shifts in polyunsaturated fatty acids (PUFA) and related metabolites after limb amputation. Linoleic acid, an omega-6 PUFA^53^, was decreased at 7dpa. The oxidized C18:2 (e.g., linoleic acid) fatty acid derivatives 9-HpODE and 9,10-DiHOME were consistently decreased at all time points. Fatty acids C20:5 (e.g., eicosapentaenoic acid, EPA) and C20:3 (e.g., eicosatrienoic acid, ETrA) were also decreased at 7dpa. EPA acts as a precursor to specialized pro-resolving mediators, including resolvins, which actively dampen inflammatory responses^54^. ETrA is a rare naturally occurring PUFA that, upon incorporation into cellular phospholipids, has been shown to suppress inflammatory nitric oxide production^55^. The reduction of multiple PUFAs and oxylipins raises the possibility that amputation leads to broad PUFA remodeling in P-CSF rather than a targeted change in a single lipid class. Whether this reflects redistribution of PUFA pools toward membrane phospholipids in the CNS, increased degradation, or a broader systemic shift in circulating lipid mediators remains an open question.

## Discussion

Individuals living with limb loss are a substantial and growing portion of the population. The systemic consequences of limb loss extend well beyond the physical absence of the limb itself^4^. Understanding how biological niches such as perilymph or CSF are affected by limb loss and how regeneration-competent organisms navigate those changes is an important and largely unexplored frontier. It is worth stating how sparse the relevant literature is for the characterization of nonmammalian perilymph-CSF. No proteomic or metabolomic characterization of perilymph or CSF exists for any salamander species. Examples of analogous works include a time-resolved proteomics study of whole otic vesicles in *Xenopus laevis* tadpoles^56^, which captures trends in inner ear development but lacks the isolation of perilymph alone. Adult avian CSF has been profiled by LC-MS in chicken and other birds to investigate adult neurogenesis^57^. Zebrafish embryonic CSF has been studied for its role in neuroepithelial survival signaling^58^, though this was not in an injury or regeneration context. The axolotl datasets reported here are therefore the first of their kind in any salamander. In combination with perturbation of regeneration informed by pathways that change in P-CSF after limb amputation, this work establishes a molecular framework for future studies aimed at determining the origins, regulation, and functional significance of P-CSF associated changes during limb regeneration.

Our work characterizing the axolotl P-CSF proteome during homeostasis revealed a diverse environment consisting of protease and inhibitor networks, lipid/carrier transport proteins, and an abundance of metabolic enzymes. This diversity is consistent with P-CSF proteins potentially being derived from multiple biological origins, including local secretory epithelium^8^, systemic circulation^13^, inner ear structures^59^ and neural tissue^60^. The presence of albumin, complement proteins, and transferrin as the most readily detected proteins is consistent with the known composition of mammalian CSF^27^. The additional recovery of inner ear associated proteins reflects the perilymph contribution to P-CSF and highlights the expected multi-sourced nature our fluid collection strategy. In our amputation timeseries, we observed a large number of decreasing proteins that progressed with limb regeneration. As no change in total abundance was observed across groups (Supplementary Fig. 1A), the apparent decreases in relative protein abundance may be indicative of a compositional redistribution of the P-CSF proteome as other proteins increase in abundance. This trend may be alternatively explained by shifting ratios of perilymph to CSF given our method of extraction. However, active repression or degradation of specific proteins cannot be ruled out.

A striking feature of the P-CSF proteomic response to limb amputation was the early and sustained elevation of sarcomeric and structural muscle proteins such as MYOM2, MYBPC2, TTN, and TCAP. These proteins were elevated as early as 3dpa and remained among the most consistently elevated proteins through 14dpa. These proteins can be detected in mammalian CSF during homeostasis, but in trace amounts^29,30^. Their coordinated elevation in axolotl P-CSF following amputation suggests that the substantial changes in muscle architecture required for limb regeneration may somehow become reflected in the P-CSF. While we considered collection contamination to be a factor in the detection of these proteins, their coordinated and sustained increase across time suggests that this may be not wholly attributable to simply contamination at the site of collection. One explanation for this finding could be that muscle undergoes a process similar to rhabdomyolysis during limb regeneration. Rhabdomyolysis is the release of intracellular muscle components from damaged or disassembled skeletal muscle into the circulation^61^. Muscle histolysis is a hallmark feature of damaged tissue near the amputation site during salamander limb regeneration^62,63^. It is therefore possible that muscle fragments may enter systemic circulation and subsequently distribute into the P-CSF. A recent study discovered that the axolotl BBB is more size permissive than that of mammals and other amphibians^64^, even during homeostasis. Amputation may further enhance BBB/BCSFB permeability, as has been observed in humans following traumatic brain injury and peripheral nerve injury^65,66^. However, these possibilities remain untested and warrant further evaluation.

We observed a transient elevation of serine protease and digestive enzymes, such as PRSS1, CTRL, CTRB2, CELA2A/3B, and CEL, at 7dpa. These enzymes are typically associated with pancreatic acinar cell secretions^67^, making their observed increase in a distant biological fluid surprising. The timing of this enrichment coincides with the early blastema stage, when cellular proliferation and ECM turnover at the amputation site are high^68^. While protease activity has known ECM remodeling roles in the regenerating blastema, less is known about how these circulating factors may affect the rest of the body. In a model of rat hemorrhagic shock, it was shown that pancreatic trypsin-, chymotrypsin-, and elastase-like enzyme activity increased in serum, peritoneal fluid, and organs distant to the digestive tract^69^. It has also been shown that direct injection of serine proteases such as porcine elastase can increase BBB permeability of mice^70^. In cluster 3 of the trajectory analysis, membership of multiple kininogen-like proteins (LOC138493245, LOC138493246, and LOC138493249) along with plasma kallikrein-like (LOC138516427) were observed. Kininogen and kallikrein-like proteins are part of the kallikrein/kinin system. Kallikreins are a subgroup of serine proteases that cleave peptides such as kininogens to form active peptides like bradykinin in humans^71^ and bradykinin-related peptides in amphibians^72^. Action of human bradykinin on one of its corresponding receptors, bradykinin receptor B2 (BDKRB2) has been shown to regulate inflammation, increase nitric oxide release, and increase BBB permeability^73^. While our data suggests an elevation of proteases in P-CSF, the origin of these proteins and consequence of their presence remains speculative.

Our data also suggests the interesting possibility that secreted proteins made in blastema cells, such as KAZALD2, SPARC, and VWDE, might reach the P-CSF. However, more experimentation would be required to test whether these proteins originate directly from the blastema or have other explanations for their appearance in P-CSF. For example, limb amputation involves nerve transection, and injured sensory nerves in the periphery could upregulate KAZALD2 expression, concomitantly elevating KAZALD2 in the P-CSF. Many other regenerative contexts in axolotls also provoke KAZALD2 upregulation such as mandible, brain, spine, and retina^74^ and, hence, testing whether these injuries also elevate KAZALD2 in P-CSF is essential for determining whether the effects we observed are specific to regenerating limbs.

The injury specificity experiments resulted in an interesting observation. The near-complete absence of differentially abundant proteins following crush injury, tail amputation, and denervation compared to the response observed after limb amputation may suggest that changes in P-CSF proteins are not simply sensitive to trauma in general. Rather, the response appears to be tied to substantial tissue loss and the formation of a blastema. Out of all the recovered proteins for both cohorts, 671 were unique to the timeseries, 254 were unique to the injury model cohort, and 1,955 proteins were shared between both. Interestingly, limb-regeneration associated proteins previously discussed such as KAZALD2 and AGR2, digestive enzymes like CEL and other trypsin-like proteins, as well as inner ear-associated protein OTOG were only recovered in the limb amputation cohort. While this may be due to natural detection variability between cohorts, their relation to limb regeneration should be noted. However, another important caveat concerns the severity of damage across injury models. In models of axolotl systemic activation^75,76^, cells throughout the body selectively re-enter the cell cycle and exhibit positive correlations with the severity of injury. Although all four limbs were injured in two of the injury models, it is possible that a certain damage threshold was not sufficient to activate the full-fold effect size observed after limb amputation. Future experiments using graded injury severity or other injuries would help to disentangle injury magnitude from injury type as the driver of our results.

The single significant hit in the denervation group, LRRN1, is interesting due to its CNS and cochlear expression profile^39^. LRRN1 is a type I transmembrane protein that plays a role as a synaptic cell adhesion molecule and tissue organizer during nervous system development^77^. It also has known roles in maintaining the pluripotency of human embryonic stem cells^78^. Its elevation in P-CSF following both nerve transection and limb amputation may reflect a local neural response to peripheral axotomy that is detectable in the P-CSF compartment. LRRN1 elevation after denervation and limb amputation may reflect a neural or otic capsule-associated response to peripheral nerve injuries, but its absence after tail amputation suggests that it is not simply a generic consequence of nerve transection. This could indicate differences in the affected nerve populations, injury anatomy, or signal access to the P-CSF compartment. This hypothesis may be worth investigating directly, particularly given the known role of innervation in blastema maintenance^79^.

The untargeted metabolomics dataset reported here is, to our knowledge, the first characterization of the axolotl P-CSF metabolome, and one of very few metabolomic profiles of CSF in any non-mammalian vertebrate. We found that the metabolites in axolotl P-CSF are consistent with the types and levels of metabolites found in mammalian CSF, such as creatinine, glutamine, and branched-chain amino acids^43^. These similarities hint at a possible overall analogous function of CSF between amphibians and mammals.

One particularly interesting metabolite we readily found in the P-CSF of uninjured axolotls is betaine. Betaine is known to act as an osmolyte and methyl donor^42^, and recent evidence shows it can act to modulate neuroinflammation and GABAergic signaling^80,81^. Betaine can suppress the NLRP3 inflammasome in mammals^82^, which can dampen neuroinflammation associated with amyloid-β. Betaine can regulate the reuptake of GABA via interaction with GABA transporter 1 (GAT1)^83^. We also detected betaine-homocysteine S-methyltransferase (LOC138511624; BHMT) in axolotl P-CSF. In mammals, this enzyme recycles homocysteine, which can be toxic, back to methionine and dimethylglycine using betaine^84^. These connections provide evidence that core aspects of axolotl P-CSF share homology with established aspects of mammalian CSF, and they suggest future lines of mechanistic experimentation to explore.

Following amputation, we observed a substantial shift in the metabolic proteins and metabolites that were detected in axolotl P-CSF. Trajectory-based analysis of P-CSF proteins during limb regeneration highlighted shifts in carbon metabolism, adenyl nucleotide binding, and amino acid biosynthesis. These themes were mirrored in the metabolomics data, suggesting the proteome and metabolome of axolotl P-CSF changes in a coordinated manner in response to limb amputation. A consistent theme across the proteomic and metabolic data was a shift in oxidative stress and antioxidant metabolism. Methionine sulfoxide (MetO) was persistently elevated across all three post-amputation time points. This metabolite often forms when ROS oxidizes methionine. Additionally, we found peroxiredoxin family members (PRDX1/2/5/6), which act as antioxidants^85^, to be decreased after amputation. Another example is PARK7, a redox-sensitive chaperone with broad neural expression^86^, which we found to decrease from baseline at both 7dpa and 14dpa. Observing a sensitive ROS marker to rise while simultaneously finding several antioxidant-associated proteins to diminish may suggest a shift in the oxidative balance of axolotl P-CSF during regeneration. This is of particularly interest because in several other models, ROS are required for regeneration^87,88^. For example, one reported role of ROS in axolotl limb regeneration is to influence blastema cell proliferation^89^.

The changes we uncovered in oxylipin and PUFA levels during regeneration are also interesting. We found decreases in linoleic acid, 9-HpODE, and 9,10-DiHOME across post-amputation time points, accompanied by decreases in EPA and ETrA at 7dpa. These data suggest axolotls may broadly remodel the PUFA landscape during regeneration. In mammals, oxylipins 9-HpODE and 9,10-DiHOME are pro-inflammatory and can have cytotoxic properties^90^, suggesting axolotls might actively dampen these metabolic pathways during regeneration. 9,10-DiHOME has also been associated with impaired leukocyte function and mitochondrial disruption^91^, so these data may also be consistent with a shift away from pro-inflammatory lipid mediator signaling. However, we also detected declining abundance of both EPA and ETrA, omega-3 PUFAs with anti-inflammatory and pro-resolving properties in mammals^55,92^. This complication may suggest that a simple shift away from inflammation and toward inflammatory resolution is unlikely to be the full story. Further investigation, including experimentation, is required to fully understand how inflammation at a systemic level may be regulated during axolotl limb regeneration and how P-CSF might participate in systems-level inflammatory processes.

The histamine catabolism signature with persistent elevation of N-acetylhistamine, ImAA, and Me-ImAA across multiple time points may suggest altered histamine metabolism and potentially increased histamine turnover. Histamine plays modulatory roles in vascular permeability^93^, immune regulation^94^, and neural activity^95^. Altered catabolism of this signaling molecule in P-CSF during regeneration may reflect changes in some or all of these processes after limb amputation. Whether this signal originates from wound-proximal immune activity, from systemic circulation via a transiently permeabilized BCSFB, or from local neural and epithelial sources cannot be resolved from the present dataset.

Another recurring theme in our metabolomics data centers on a potential role for an arginine-proline axis. We found the arginine catabolite 4-Guanidinobutyric acid (4-GBA)^96^ to be enriched at 7 and 14dpa. This elevation was accompanied by elevated levels of proline and citrulline at 7dpa and ornithine at 14dpa. One possible explanation for an elevated arginine-to-proline substrate chain could be the increased demand for collagen biosynthesis during limb regeneration since proline is a crucial component of collagen^97^. This signature coincides with the peak ECM enrichment we observed in the proteomics dataset. Separately, 4-GBA has been identified as a neuroactive compound that functions as a mimetic of γ-aminobutyric acid (GABA) and agonist of GABA-A receptors^98^. Intracranial treatment with muscimol, a GABA agonist, has recently been shown to decrease the rate of axolotl tail regeneration by decreasing the excitability of glutamatergic neurons in the telencephalon^99^. A speculative, but mechanistically interesting possibility is that arginine catabolism during regeneration generates metabolites that influence GABA’s inhibitory activity in regions with access to P-CSF. Such a hypothesis requires functional testing, but developing it, and many more, illustrates the value of an untargeted metabolomics approach to reveal connections between known regenerative processes and neural signaling chemistry.

This study demonstrates that axolotl P-CSF undergoes coordinated compositional changes following limb amputation. The specificity of these changes to limb amputation rather than other injury types, the consistency of cross-omics themes around oxidative stress, ECM remodeling, lipid mediator dynamics, and the detection of established blastema-associated proteins in this fluid collectively suggests that P-CSF may serve as a record of systemic responses to limb loss and as a potential conduit through which neural and circulating signals interface with the regenerating tissue. However, much remains to be determined about the fluid’s cellular sources, barrier dynamics, and functional contributions. This study opens a new analytical vantage point on one of biology’s most remarkable phenomena, and we hope it serves as a resource and a provocation for the field.

## Methods

### Animals

All animal experimentation was approved by and conducted in accordance with Harvard University’s Institutional Animal Care and Use Committee. Wild type leucistic axolotls (*Ambystoma mexicanum*) were housed individually to ensure they remain naïve to bite injury before experimentation. Animals were housed in 2.63 g/L Instant Ocean water, 4000 μS conductivity. The animals were kept at 18°C with a 14 h light/10 h dark cycle and were fed axolotl sinking soft moist pellets (Aquatic Foods). Age-, sex, and size-matched (10-18 months old, 12-18cm tail-to-snout length) animals were randomly assigned to each experimental group. Cohorts were single-sex and chosen based on animal availability at time of collection. Sex was determined using examination of physical sexual characteristics. For P-CSF blood comparisons, all females were used. All females were used for the proteomics cohorts, and all males were used for the metabolomics cohort. The number of replicates used in each experiment refers to individual animals unless otherwise noted.

### Amputation, denervation, and crush injuries

Animals in all experiments were size, sex, and age matched. Animals were anesthetized in 0.1% (w/v) tricaine prior to all procedures involving control groups, amputation, denervation, or crush injury. After all surgical procedures, axolotls were allowed to recover overnight in 0.5% (w/v) sulfamerazine. All limb amputations were performed at the mid-stylopod for all four limbs. The bone was then trimmed back from the amputation plane. For forelimb denervations, the brachial plexus nerves of both forelimbs were transected according to Schotté *et al*^100^ and for hindlimb denervation, the sciatic nerves were transected according to Kropf *et al*^101^. Incisions were made in the axial regions corresponding to the brachial plexus (forelimb) and sciatic nerves (hindlimb). Exposed nerves were then isolated with forceps and a small section was resected (∼1mm) using fine scissors to achieve complete denervation of both forelimbs and hindlimbs. For crush injuries, straight forceps were used to apply pressure to the mid-stylopod for 30 seconds as described in^76^ of all four limbs. For tail amputations, 2cm from the tail tip was fully amputated using fine scissors.

### P-CSF extraction

All animals were anesthetized as mentioned above prior to P-CSF extraction. An “L-shaped” incision was made on the dorsal side of the head. Skin was peeled back on one side to reveal muscle. Small scissors were used to carefully remove muscle tissue away from the skull, revealing the otic capsule. A Dremel tool was used to carefully debride excess tissue and thin the bone at the extraction site. A pulled 150mm borosilicate glass capillary needle attached to a 1mL syringe was then used to pierce the otic capsule. A clear, colorless fluid should begin to slowly flow through the needle. Negative pressure was periodically applied to the syringe plunger if fluid flow stopped. Approximately 80-90µL of clear P-CSF was extracted per needle volume per animal over the course of 1 hour. Amphibian erythrocytes are nucleated and are up to ∼6-fold larger in diameter than human erythrocytes^102^, simplifying preliminary visual inspection of blood contamination. Any samples with visible blood contamination were discarded. Immediately following P-CSF extraction, samples were placed in LoBind Eppendorf tubes on ice. Samples were then spun at 2,000g for 10min at 4°C. Any samples with red blood cell pellets were discarded. Samples were then aliquoted and stored at –80°C until further analysis.

### Fluorescently labeled 10-kDa dextran injection

For dye tracing experiments, animals were prepped similarly to the P-CSF extraction procedure as previously mentioned. 10 µl of 10-kDa dextran conjugated with tetramethylrhodamine (Invitrogen) was injected directly into the otic capsule using a pulled glass capillary. This molecular weight was selected to be a tracer that is larger than what can typically passively diffuse through an intact mammalian BCSFB while remaining small enough to flow freely through open fluid connections^103^. Animals were then imaged using a stereomicroscope (Leica Microsystems) using the brightfield and RFP filter set.

### P-CSF blood comparisons

Cell count of intact P-CSF and whole blood was calculated manually using 10 µL of either P-CSF without centrifugation or whole blood diluted 1:200 in 0.7X PBS using a hemocytometer (n = 6). Total protein concentration of intact P-CSF and plasma was quantified using BCA assay (Thermo Scientific) (n = 6). 5 µL of sample were used per reaction and ran in triplicate. Glucose concentration of intact P-CSF and whole blood was determined using a glucose meter (CVS Health) (n = 6). 5 µL of sample was used per test. Each P-CSF sample was tested for the presence of hemoglobin using a Hemoglobin High Sensitivity Calorimetric Detection Kit (Arbor Assays). 5 µL of sample were used per reaction and ran in duplicate. Any samples that contained more than 1.5 µg/mL hemoglobin were discarded per the threshold recommended by Paciotti et al^104^. Statistical comparisons were made using a two-sided unpaired t-test with Welch’s correction.

### Proteomics sample preparation

16 P-CSF samples (n = 4 each group) were chosen to submit for the amputation timeseries proteomic analysis. For the injury model proteomics, 14 P-CSF samples were submitted (n = 4 each for Intact and Crush; n = 3 each for Tail Amp and Denervation). Each sample was tested for total protein and hemoglobin concentration as previously mentioned. Statistical comparisons were made using a two-sided Dunnett’s T3 multiple comparisons test.

Equal protein amounts from each axolotl CSF sample were aliquoted into a new protein low binding microcentrifuge tube and adjusted to a final volume of 100 µL with 100 mM triethylammonium bicarbonate (TEAB). Proteins were reduced with 10 mM tris(2-carboxyethyl) phosphine (TCEP) at 55 °C for 1 hour and then alkylated with 15 mM iodoacetamide in the dark for 30 minutes. Proteins were then precipitated using chloroform-methanol precipitation method and resuspended in 100 mM TEAB. Samples were digested with a Trypsin/Lys-C mix overnight at 37 °C.

### TMTpro 16plex labeling

TMTpro labeling was performed according to manufacturer’s instructions (Thermo Fisher Scientific). Briefly, TMTpro labeling reagents were dissolved in 20 µL of anhydrous acetonitrile and then added to samples in equal volume. After one hour incubation at room temperature, the reaction was quenched with 5 µL of 5% hydroxylamine. Labeled peptides were then combined in equal amounts and dried in a SpeedVac.

### High pH reverse phase fractionation

TMT-labeled peptides were fractionated using an Agilent 1200 HPLC system (Santa Clara, CA) using XBridge BEH C18 column, 250 x 4.6 mm, 3.5 μm with XBridge BEH C18 VanGuard Cartridge 3.9 mm x 5 mm, 3.5 μm (Waters, Milford, MA). The mobile phases used for separation were 10 mM ammonium formate in water (mobile phase A) and 10 mM ammonium formate in 90% acetonitrile/10% water (mobile phase B). Peptides were separated using a 76 min gradient, from 0 % to 35% mobile phase B in 66 min and increased to 70% mobile phase B in 10min. Elution was collected into 20 fractions and then dried completely in a SpeedVac (Eppendorf, Germany). Fractions were resuspended in 0.1% formic acid in water before LC-MS/MS analysis.

### Proteomics LC-MS/MS

Samples were analyzed on an Orbitrap Eclipse Tribrid Mass Spectrometer coupled with Vanquish Neo UHPLC system (Thermo Fischer, Waltham, MA) at Harvard Center for Mass Spectrometry. TMT labeled peptides were first trapped on a trapping cartridge (300µm x 5mm PepMap™ Neo C18 Trap Cartridge, Thermo scientific) prior to separation on an analytical column (µPAC, C18 pillar surface, 50 cm bed, Thermo scientific) with column temperature at 35 °C. The mobile phase system consisted of water with 0.1% FA (A) and acetonitrile with 0.1% FA (B). Peptides were separated with a gradient elution from 2 – 40% B over 180 minutes with a flow rate of 350 nL/min. The mass spectrometer operated in data-dependent acquisition (DDA) mode for all analyses. Electrospray positive ionization was enabled with a voltage at 2.1 kV. FAIMS compensation voltages of –40, –55, –70 V were combined in a single run with a cycle time of 1.5 s each. Full scan ranging from 400 – 1600 m/z was performed with a mass resolution of 12×10^4^ and with AGC target set to Standard and maximum injection time set to Auto. The MS2 was operated with resolution 5.0×10^4^ and AGC target was 250%. Maximum injection time was 200 ms and HCD collision energy of 38%.

### Proteome Discoverer

Raw data was submitted for analysis in Proteome Discoverer 3.0 software (Thermo Scientific). The MS/MS data was annotated against the latest axolotl genome assembly UKY_AmexF1_1 (RefSeq assembly: GCF_040938575.1) along with known contaminants such as human keratins and common lab contaminants. Sequest HT searches were performed using the following guidelines: a 10 ppm MS tolerance and 0.02 Da MS/MS tolerance; Trypsin digestion with up to two missed cleavages; carbamidomethylation (57.021 Da) on cysteine and TMTpro 16-plex tags on both peptide N-termini and lysine residue (+304.207 Da) were set as static modification; oxidation (+15.995 Da) of methionine was set as a variable modification; Minimum required peptide length set to ≥ 6 amino acids. At least one unique peptide per protein group is required for identifying proteins. All MS2 spectra assignment FDR of 0.01 on both protein and peptide level was achieved by applying the target-decoy database search by Percolator. Reporter ion abundances were normalized within Proteome Discoverer using the total peptide amount normalization mode parameter. A per-channel scaling factor was calculated from summed peptide-group abundances such that total abundance was equalized across all TMT channels.

### Gene annotation

The axolotl genome annotation assigns placeholder identifiers (e.g., LOCXXXXXXXXX) to uncharacterized or computationally predicted loci^25^. For downstream analyses, LOC-annotated entries were assigned gene symbols using NCBI gene descriptions. Entries with gene descriptions were assigned to the closest matching human ortholog where applicable. Proteins sharing the same gene symbol were distinguished by sequentially appending a numeric suffix (e.g., GENE_1, GENE_2). Any genes that could not be assigned gene symbols retained their LOC-identifier. As a result, all proteins with no matching human symbol were excluded from downstream pathway analyses. Each protein was assigned a protein class using the PANTHER classification system^26^ and then manually consolidated into generalized categories (Supplementary Table 2). Proteins with no PANTHER classification were labeled “Unclassified”.

### Proteomics statistical comparisons

Proteomic data analysis was performed in R (v4.4.2). Raw normalized TMT abundances were exported from Proteome Discoverer and imported for reanalysis; pre-computed abundance ratios and p-values provided by Proteome Discoverer were not used for statistical inference. Normalized abundance values were log2-transformed, and proteins present in fewer than 75% of replicates within all groups were removed prior to analysis. Differential abundance was assessed using linear models fitted with the limma package^105^, followed by empirical Bayes variance moderation via eBayes(), which stabilizes per-protein variance estimates by borrowing information across the full detected proteome. Peptide-count-weighted variance moderation was then applied using the DEqMS package^106^, which further adjusts each protein’s variance prior according to the number of unique quantified peptides.

For the limb amputation timeseries, three pairwise contrasts were tested against the Intact reference condition (3 dpa vs. Intact, 7 dpa vs. Intact, and 14 dpa vs. Intact). One outlier per group was removed based on PCA clustering for a final sample size of n = 3 per group). PCA without outlier removal is available in (Supplementary Fig. 1E). For the injury comparison dataset, three contrasts were tested at 14 days post-injury (Denervated vs. Intact, Crush vs. Intact, and Tail amputation vs. Intact). One outlier was removed from the Intact and Crush groups for a final sample size of n = 3 per group. PCA without outlier removal is available in (Supplementary Fig. 2D). Statistical significance was determined using permutation-based false discovery proportion (permFDP) estimation^107^ applied separately to each contrast using the DEqMS-moderated p-values (sca.P.Value). This approach derives a contrast-specific corrected p-value rejection threshold controlling the expected false discovery proportion across permutations of the data. 1,000 permutations were used in order to exceed the maximum number of distinct permutations. Benjamini-Hochberg false discovery rate-adjusted p-values (sca.adj.pval) are reported alongside all results for transparency but were not used as the primary criterion for significance calling. All statistical analyses were performed using R/Bioconductor, with figures generated using ggplot2.

### Gene Ontology, KEGG, and PCA

Gene Ontology (Biological Process) and KEGG pathway enrichment analyses were performed in R using the clusterProfiler package(v4.14.16)^108^. Since axolotls are not fully represented in gene ontology and KEGG databases, enrichment analyses were performed using the human genome database (org.Hs.eg.db) as a reference. For each group analyzed, enrichment was tested against a background gene universe restricted to all proteins detected within the dataset. Both analyses used Benjamini-Hochberg-adjusted p-value and q-value cutoffs of 0.05 and 0.25, respectively. PCA was performed for the limb amputation timeseries and injury-comparison datasets using the log2-transformed abundance values. PCA was computed using the prcomp() function in R with unit-variance scaling applied to each protein.

### Trajectory clustering analysis

To identify temporal patterns of protein abundance across the limb amputation timeseries, proteins were subjected to fuzzy c-means clustering using the Mfuzz package^109^ in R. For each protein, mean log2 abundance was calculated within each timepoint and the resulting trajectory was standardized to a z-score (mean = 0, SD = 1) across timepoints. Fuzzy c-means clustering was performed using the mfuzz() function with the fuzzifier parameter set to m = 1.25. The number of clusters (k = 6) was selected by evaluating the elbow plot of minimum pairwise inter-centroid distances (Supplementary Fig. 3). All proteins with scores exceeding 0.55 were assigned to the cluster for which it had the highest membership score. Remaining proteins were listed as “Unassigned”.

### Metabolomics sample preparation

P-CSF samples were prepared for untargeted metabolomics at the Harvard Center for Mass Spectrometry. A total of 20 samples (n = 5 each group) were prepared for initial submission. Briefly, 100 µL of methanol was dispensed into low-absorption 1.5 mL microcentrifuge tubes, and 25 µL of each P-CSF sample was transferred into the methanol for protein precipitation. Samples were centrifuged at –11°C for 2 hours. Supernatants were transferred to new tubes and dried under nitrogen flow. Dried extracts were resuspended in 20 µL of 30% acetonitrile in water, and 14 µL of each sample was transferred to glass microinserts for injection. The remaining volume from all samples was pooled to generate a pool sample used for AcquireX deep scan acquisition.

### Metabolomics LC-MS/MS

Samples were analyzed on a Thermo ID-X Tribrid mass spectrometer coupled with a Vanquish UHPLC system (Thermo Fisher Scientific) at the Harvard Center for Mass Spectrometry. Metabolites were separated on a Waters Atlantis Premier BEH Z-HILIC column (1.7 µm, 2.1 × 150 mm) with a Vanguard Fit guard column, maintained at 40°C with samples held at 4°C. An injection volume of 5 µL was used. Mobile phase A consisted of 20 mM ammonium carbonate with 0.1% ammonium hydroxide in water, and mobile phase B consisted of 97% acetonitrile in water. Separation was performed using a 45-minute gradient starting at 93% B, decreasing to 40% B at 19 minutes, reaching 0% B at 28 minutes, holding until 33 minutes, and returning to 93% B by 36 minutes. The flow rate was 0.05 mL/min initially, increasing to 0.15 mL/min. The column eluate was diverted to waste from 0–1 minutes and 32–45 minutes. Data were acquired in data-independent mode using alternating positive and negative ionization (HESI) with a resolution of 240,000, RF lens 30%, normalized AGC target 25%, maximum injection time 50 ms, and a scan range of 65–1,000 m/z. The pooled sample was analyzed using AcquireX deep scan MS2/MS3 acquisition at five levels in both negative and positive modes independently to build a compound-specific spectral library for the dataset.

### Metabolomics data processing

The resulting raw LC-MS files, including sample files (MS1), identification files (MS2/MS3), and blank runs, were analyzed by Compound Discoverer (Thermo Fisher Scientific, version 3.3) using default metabolomics workflow including online database (mzCloud, Mass List, and ChemSpider) searching. Compounds with a minimum of peak shape score 3 in 80% of sample files were included for analyses. The resulting compound table was filtered as follows: **(1)** compound is not background, **(2)** compound name is not blank; **(3)** annotation ΔMass is within 5 ppm; **(4)** compound has full match in Predicted Compositions; **(5)** compound has full match in mzCloud search (MS2 spectra required). Next, the compound list was manually curated, including removing features with bad peak shapes and features not likely of biological origin. “Full-match” and “partial match” mzCloud-based annotations that did not appear as the top-ranked matches were also retained in the data sheet to represent additional possible annotations. Chemical class annotations were assigned to each feature using a curated HMDB-ID-based lookup table. Features that could not be assigned by either method were designated as “Unclassified”.

### Metabolomics statistical comparisons

Metabolomic data analysis was performed in R (v4.4.2). Normalized peak areas exported from Compound Discoverer 3.3 were imported for downstream processing. Prior to statistical analysis, sample outliers were identified and removed, resulting in a final sample set of (n = 3) per time point. Differential abundance was assessed using linear models fitted with the limma package, followed by empirical Bayes variance moderation via eBayes() with trend and robust estimation enabled to stabilize variance estimates across features. Three pairwise contrasts were tested against the Intact reference condition (3 dpa vs. Intact, 7 dpa vs. Intact, and 14 dpa vs. Intact). Statistical significance was determined using permFDP estimation, applied separately to each contrast using the limma-moderated p-values as described previously. Benjamini-Hochberg FDR-adjusted p-values are reported alongside all results for transparency but were not used as the primary significance criterion. All statistical analyses were performed using R/Bioconductor, with figures generated using ggplot2.

### Use of generative AI and AI-assisted technologies

Claude Sonnet 4.6 was used to streamline R-based data analysis pipeline generation as well as improve readability and conciseness of early manuscript drafts.

## Data availability

The raw mass spectrometry proteomics data generated in this study will be deposited in the PRIDE partner repository (ProteomeXchange Consortium) prior to publication. The metabolomics data generated in this study will be deposited in Metabolomics Workbench (National Metabolomics Data Repository, NMDA) prior to publication.

## Code availability

Custom analysis code will be made publicly available via a Zenodo-archived GitHub repository prior to publication.

## Acknowledgements

The authors thank Mei Chen, Steven Kolakowski, and Charles Vidoudez at the Harvard Center for Mass Spectrometry for their assistance in sample preparation, mass spectrometry data acquisition, and data analysis. This material is based upon work supported by the National Science Foundation Graduate Research Fellowship under Grant No (2140743). Any opinion, findings, and conclusions or recommendations expressed in this material are those of the authors(s) and do not necessarily reflect the views of the National Science Foundation. Additional support was provided by the Harvard Faculty of Arts and Sciences Dean’s Competitive Fund, the HCBI Simmons Award, and Harvard University. E.T.C. is an HHMI investigator. We would also like to thank Steven P. Gygi for his guidance and support on this work.

## Author contributions

Conceptualization, N.L. and J.L.W.; data curation, N.L., B.Z., and S.R.S.; formal analysis, N.L., B.Z., and S.R.S.; funding acquisition, N.L. and J.L.W.; investigation, N.L., B.Z., S.R.S., Y.Z., D.P., J.C.P., S.Y.C.W., T.P., K.C., S.B., H.D.S., A.R.J., R.T.K., L.S., E.T.C., and J.L.W.; methodology, N.L., B.Z., and S.R.S.; project administration, N.L., H.D.S. and J.L.W.; resources, N.L. and H.D.S.; software, N.L., B.Z., and S.R.S. and H.D.S.; supervision, E.T.C. and J.L.W; validation, N.L.; visualization, N.L.; writing – original draft, N.L.; writing – review & editing, N.L., B.Z., and S.R.S., D.P., J.C.P., S.Y.C.W., H.D.S., R.T.K., E.T.C., and J.L.W.

## Figure Legends

**Supplementary Figure 1:**
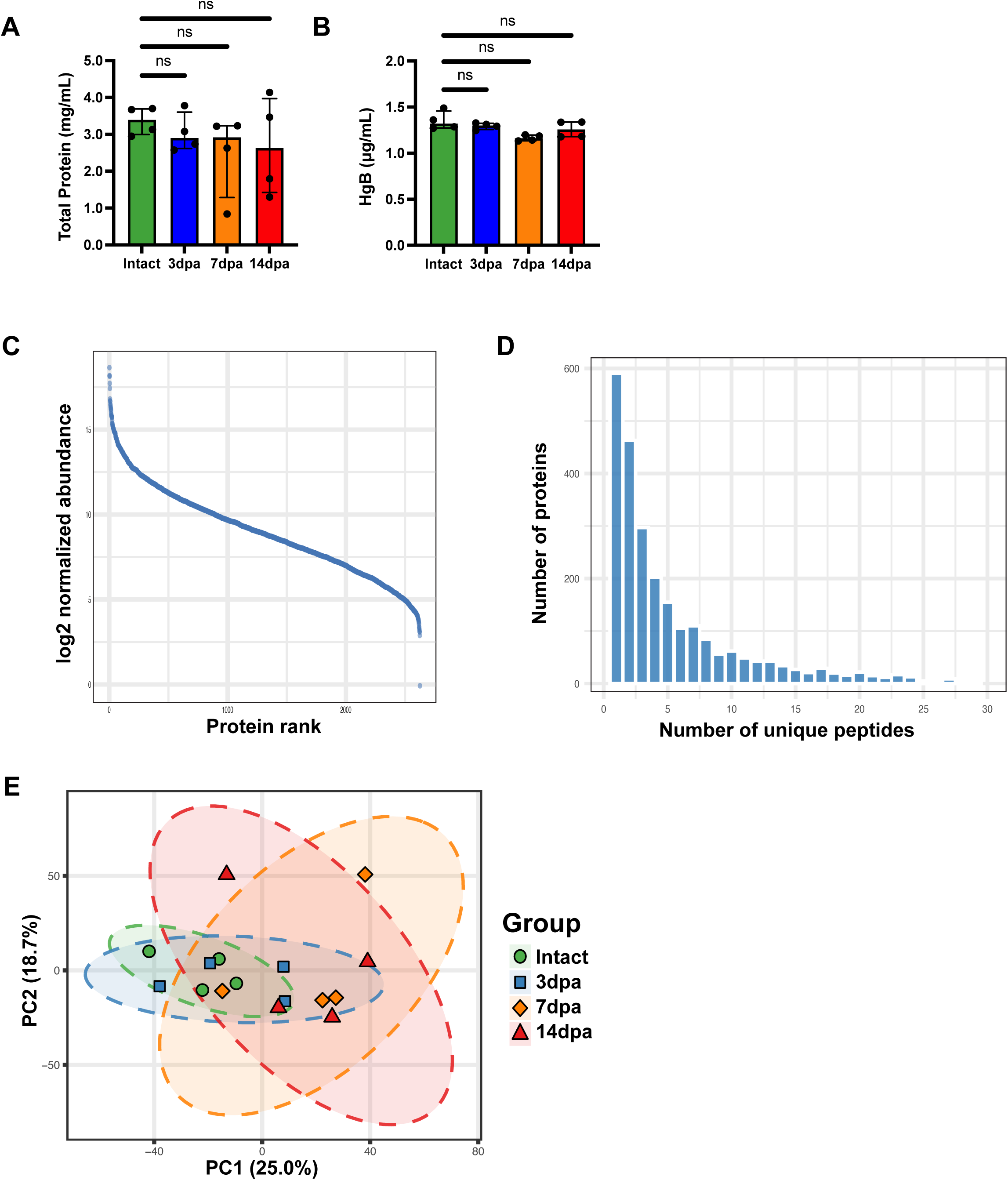
Quality control for P-CSF limb amputation proteomic samples. **a** BCA mean total protein concentration of P-CSF samples from all submitted limb amputation time points. Two-sided Dunnett’s T3 multiple comparisons test. Intact vs 3dpa (adj p-value = 0.7129). Intact vs 7dpa (adj p-value = 0.4560). Intact vs 14dpa (adj p-value = 0.7185). Data are presented as mean ± s.d. ns: not significant. **b** Mean hemoglobin concentration of P-CSF samples from all submitted limb amputation samples. Two-sided Dunnett’s T3 multiple comparisons test. Intact vs 3dpa (adj p-value = 0.6657). Intact vs 7dpa (adj p-value = 0.0587). Intact vs 14dpa (adj p-value = 0.4797). Data are presented as mean ± s.d. ns: not significant. **c** Dynamic range plot of normalized relative protein abundance values across all samples. Protein rank is represented on the x-axis. log2 normalized protein abundance is represented on the y-axis. **d** Histogram of unique peptides across identified proteins. Number of unique peptides is represented on the x-axis. Number of proteins is represented on the y-axis. **e** PCA plot of P-CSF limb amputation timepoints prior to outlier removal.

**Supplementary Figure 2:**
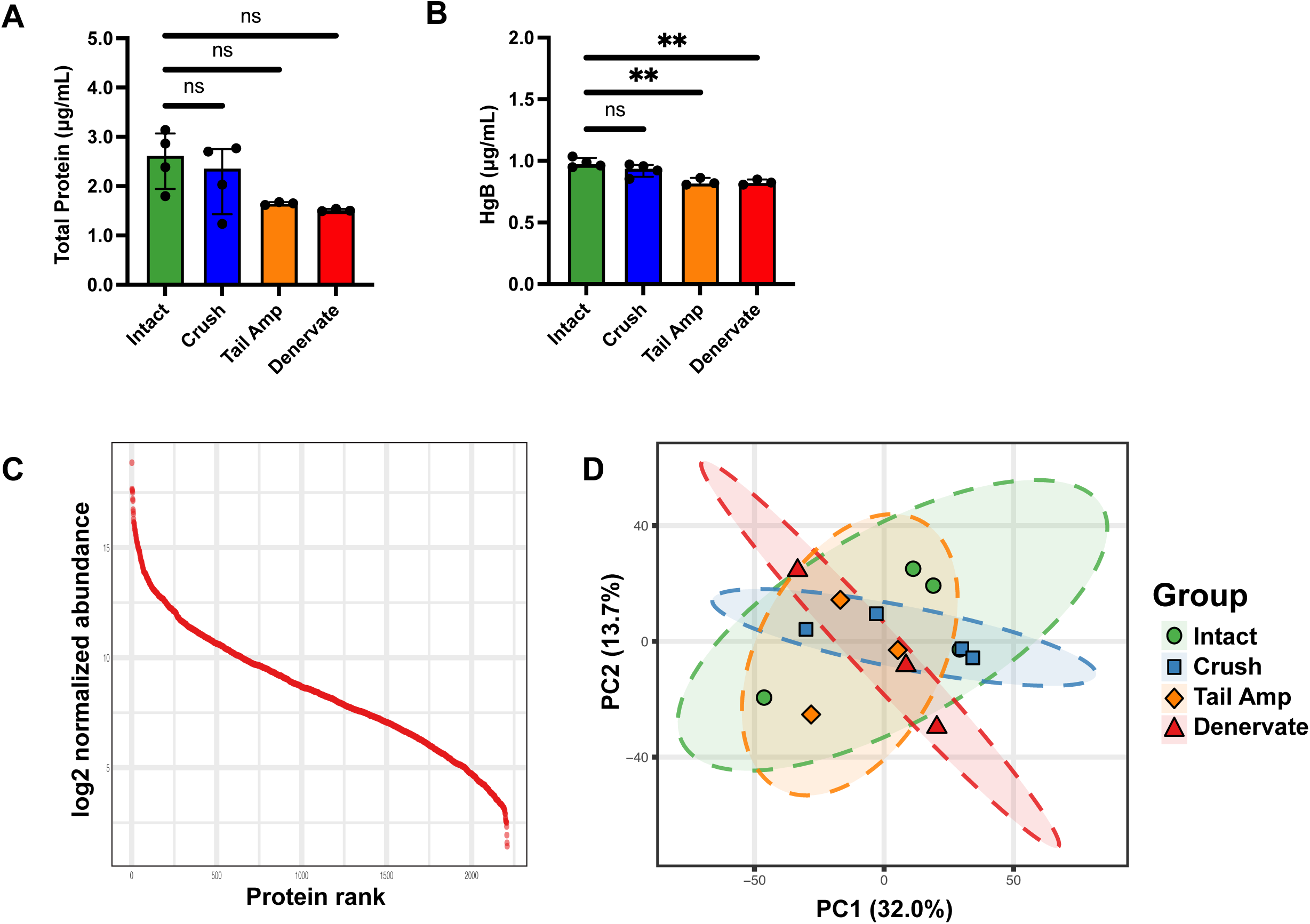
Quality control for P-CSF injury model proteomic samples. **a** BCA mean total protein concentration of P-CSF samples from all submitted injury model samples. Two-sided Dunnett’s T3 multiple comparisons test. Intact vs Crush (adj p-value = 0.8190). Intact vs Tail Amputation (adj p-value = 0.1266). Intact vs Denervation (adj p-value = 0.0902). Data are presented as mean ± s.d. ns: not significant. **b** Mean hemoglobin concentration of P-CSF samples from all submitted injury model samples. Two-sided Dunnett’s T3 multiple comparisons test. Intact vs Crush (adj p-value = 0.3122). Intact vs Tail Amputation (adj p-value = 0.0025). Intact vs Denervation (adj p-value = 0.0047). Data are presented as mean ± s.d. ns: not significant. **: adj p-value < 0.01. **c** Dynamic range plot of normalized relative protein abundance values across all samples. Protein rank is represented on the x-axis. log2 normalized protein abundance is represented on the y-axis. **d** PCA plot of P-CSF injury model samples prior to outlier removal.

**Supplementary Figure 3:**
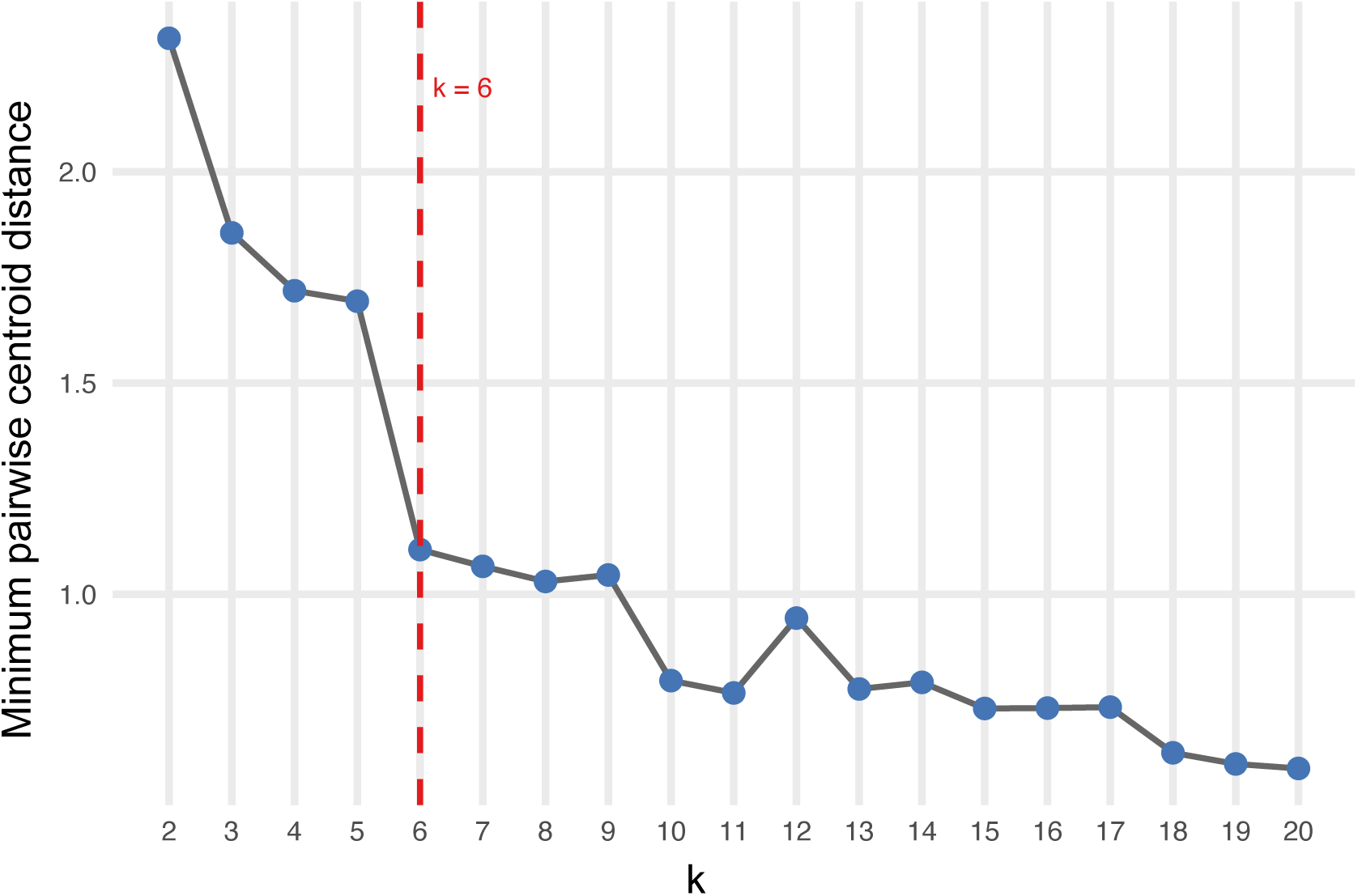
Minimum pairwise centroid distance across k-values for trajectory clustering. Graph of minimum pairwise centroid distance between cluster centroids plotted against number of clusters (k). Vertical dotted red line at k = 6 represents the selected number of clusters.

**Supplementary Table 1: Limb amputation timeseries proteomics results**

Table listing all proteins quantified across the limb amputation regeneration timeseries.

**Supplementary Table 2: PANTHER protein class reference**

Table listing PANTHER protein class terms and their corresponding collapsed classification category.

**Supplementary Table 3: Limb amputation timeseries proteomics PCA loadings**

Table listing protein loadings for principal components 1 and 2 of the limb amputation timeseries proteomics PCA.

**Supplementary Table 4: Limb amputation timeseries proteomics enriched GO terms**

Table listing Gene Ontology (GO) enrichment terms for proteins differentially abundant at each timepoint contrast.

**Supplementary Table 5: Limb amputation timeseries proteomics enriched KEGG terms**

Table listing KEGG enrichment terms for proteins differentially abundant at each timepoint contrast.

**Supplementary Table 6: Limb amputation timeseries proteomics trajectory cluster membership**

Table listing trajectory cluster assignment for each protein.

**Supplementary Table 7: Limb amputation timeseries proteomics trajectory cluster enriched GO terms**

Table listing Gene Ontology (GO) enrichment terms for each trajectory cluster

**Supplementary Table 8: Limb amputation timeseries proteomics trajectory cluster enriched KEGG terms**

Table listing KEGG enrichment terms for each trajectory cluster

**Supplementary Table 9: Injury model proteomics results**

Table listing all proteins quantified across injury models.

**Supplementary Table 10: Injury model proteomics PCA loadings**

Table listing protein loadings for principal components 1 and 2 of the injury model proteomics PCA.

**Supplementary Table 11: Limb amputation timeseries metabolomics results**

Table listing all metabolites quantified across the limb amputation regeneration timeseries.

**Supplementary Table 12: Limb amputation timeseries metabolomics PCA loadings**

Table listing metabolite loadings for principal components 1 and 2 of the limb amputation timeseries metabolomics PCA.

